# CYP1B1 converts procarcinogens into genotoxins in *Saccharomyces cerevisiae*

**DOI:** 10.1101/2021.07.22.453414

**Authors:** Akaash Kannan, Nicholas Perpetua, Michael Dolan, Michael Fasullo

## Abstract

CYP1B1 activates many chemical carcinogens into potent genotoxins, and allelic variants are risk factors in lung, breast, and prostate cancer. However, genetic instability phenotypes incurred by CYP1B1-activated metabolites have been investigated for only few compounds. In this study, we expressed human CYP1B1 in yeast strains that measure DNA damage-associated toxicity and frequencies of chromosomal translocations and mutations. DNA damage-associated toxicity was measured in a *rad4 rad51* strain, defective in both DNA excision and recombinational repair. Frequencies of chromosomal translocations were measured in diploid yeast strains containing two *his3* fragments, and mutation frequencies were measured by selecting for canavanine resistance (Can^R^) in haploid strains. These strains were exposed to benzo[a]pyrene dihydrodiol (BaP-DHD), aflatoxin B1 (AFB_1_), and the heterocyclic aromatic amines, 2-amino-3,8-dimethylimidazo[4,5-f]quinoxaline (MeIQx) and 2-amino-3-methylimidazo(4,5-*f*)quinoline (IQ). We observed that AFB_1_, BaP-DHD, IQ, and MeIQx conferred toxicity in the DNA repair mutant expressing CYP1B1. Translocation frequencies increased eight-fold and three-fold after exposure to 50 μM AFB_1_ and 33 μM BaP-DHD respectively. Only a two-fold increase in mutation frequency was observed after exposure to 50 μM AFB_1_. However, a robust DNA damage response was observed after AFB_1_ exposure, as measured by the induction of the small subunit of ribonucleotide reductase, Rnr3. While CYP1B1-mediated activation of BaP-DHD and heterocyclic aromatic amines was expected, strong activation of AFB_1_ was not. These studies demonstrate that CYP1B1-mediated activation of carcinogens does not only activate compounds to become mutagens but also can convert compounds to become potent recombinagens.

## INTRODUCTION

Many environmental carcinogens are not genotoxic *per se* and require phase I cytochrome P450 (CYP) enzymes for bioactivation. These compounds, referred to as procarcinogens, include polyaromatic hydrocarbons (PAHs) and heterocyclic aromatic amines (HAAs), which are activated to form highly reactive epoxide and nitrenium ion derivatives, respectively (Turesky and Marchand, 2011; Reed et al., 2018). Among enzymes encoded by 57 human CYP genes, CYPs 1A1, 1A2, 1B1, 2A6, 2A13, 2E1, and 3A4 function in activating carcinogens (for review, see Manikandan and Nagini, 2018; Barnes 2018; Guengerich, 2019). While hepatic CYPs are known to bioactivate many carcinogens, the extrahepatic CYPs, CYP1A1 and CYP1B1 (Ding and Kaminsky, 2003), also activate carcinogens associated with lung, bladder, breast, and skin cancer (for reviews see Koual et al., 2020).

Human cDNA encoding CYP1B1 was first cloned from dioxin-exposed keratinocyte tissue (Sutter et al., 1994) and its genomic copy, containing three exons and two introns, maps to human chromosome 2 (Tang et al., 1996). While low amounts of CYP1B1 mRNA can be detected in the liver (Chang et al., 2001), the CYP1B1 protein is primarily expressed in extrahepatic tissues, including breast, prostate, ovary, kidney and lungs (Tang et al., 1996, Spink et al., 1998; Spivack et al., 2001). Due to its role in metabolizing steroid hormones and fatty acids that stimulate growth of cancer cells (Badawi, 2001; Alsubait, 2020), CYP1B1 has been a target for therapeutic inhibition (Li et al., 2017). Particular Cyp1B1 allelic variants, which confer decreased enzymatic activity, are highly penetrant in conferring pediatric glaucoma (Falero-Perez et al., 2019), indicating a role in ocular development. CYP1B1 over-expression is also correlated with higher levels of oxidative stress and has been observed in particular tumors (Kwon et al., 2016). These studies underscore CYP1B1’s function in modulating cell growth and development.

Abundant evidence, derived from epidemiological human studies, cell cultures, and transgenic mice, suggests that CYP1B1 has a prominent role in activating a wide variety of procarcinogens (Shimada et al., 1996). CYP1B1 knock-out mice are more resistant to the carcinogenic effects of polyaromatic hydrocarbons, such as 7, 12-dimethylbenz[a]anthracene (DMBA, Buters et al. 1999), while CYP1B1 humanized mice have a higher incidence of dibenzo[def,p]chrysene-associated cancers (Madeen et al., 2017). Human CYP1B1 polymorphisms have been correlated with the higher incidence of endometrial, colon, lung and laryngeal cancer (Trubicka et al., 2010; Chen et al., 2015; Elfaki et al., 2018); among Asians, the CYP1B1 G119T allele is a risk factor for prostate and breast cancer (Li et al., 2015).

Hormone replacement therapy, which induces CYP1B1 expression, has been correlated to increase risk of cancer in women who smoke (Roos and Bolt, 2005). Pharmacological inhibition of CYP1B1 reduces the activation of carcinogenic agents in vitro (Cui and Li, 2015). When expressed in either leukocytes or in vitro, CYP1B1 has been observed to activate polyaromatic hydrocarbons (Crespi et al., 1997). These studies suggest that CYP1B1-mediated carcinogen activation could engender multiple genetic instability phenotypes; however, few genetic instability endpoints have been directly ascribed to CYP1B1-mediated activation of genotoxins.

Because *Saccharomyces cerevisiae* (budding yeast) does not contain CYPs that metabolically activate many carcinogens (for review, see Eki, 2018), “humanized” budding yeast strains have been useful in determining which human CYPs metabolically activate compounds to form genotoxins. Previous studies have shown that expression of CYP1A1 (Sengstag et al., 1996; Fasullo et al., 2017), CYP3A4 (Fasullo et al., 2017), and CYP1A2 (Sengstag et al., 1996, Fasullo et al., 2014) can activate a wide variety of carcinogens in yeast using a multitude of genotoxic endpoints, including DNA adducts, mutations, recombination, and checkpoint activation. Shimada et al. (1999) positioned the CYP1B1 cDNA under the control of the inducible yeast GAL promoter and introduced the CYP1B1 expression plasmid into budding yeast. Subsequently, they obtained cytoplasmic lysates and detected ethoxyresorufin O-deethylase (EROD) and polyaromatic hydrocarbon hydroxylation activities (Shimada et al., 1999). In this manuscript, we show that expression of CYP1B1 in yeast can activate polyaromatic hydrocarbons into potent genotoxins, increasing frequencies of carcinogen-associated translocations and mutations.

## MATERIALS AND METHODS

### Strains, Media, Chemicals

Standard media were used for the culture of yeast and bacterial strains (Burke et al. 2000). LB-AMP (Luria broth containing 100 μg/ml ampicillin) was used for the culture of the bacterial strain DH1 strains containing the vector pGAC24, pGAC24-CYP1B1, and pYES2-CYP1B1. Media used for the culture of yeast cells included YPD (yeast extract, peptone, dextrose), YP-raffinose (yeast extract, peptone, raffinose), YP-galactose (yeast extract, peptone, galactose), SC (synthetic complete, dextrose), SC-HIS (SC lacking histidine), SC-URA (SC lacking uracil), SC-LEU (SC lacking leucine) and SC-ARG (SC-lacking arginine). Media to select for canavanine resistance contained SC-ARG (synthetic complete lacking arginine) and 60 μg/mL canavanine (CAN) sulfate.

Strains to measure growth, translocation frequencies and mutations are listed in Table I. The *rad4 rad51* strains are defective in both nucleotide excision repair (NER) and recombinational repair and were used to measure growth inhibition resulting from DNA damaging agents. Strains to measure carcinogen-associated frequencies of translocations are diploid strains containing one copy of the *his3* fragments at chromosomes II and IV at the *GAL1* and *trp1* loci (Fasullo and Davis, 1988), respectively. Galactose induction of CYP1B1 was performed in Gal^+^ strains containing the *trp1-Δ69* allele. The haploid strain YB204 was used to measure frequencies of canavanine (CAN) resistance.

**TABLE I.**
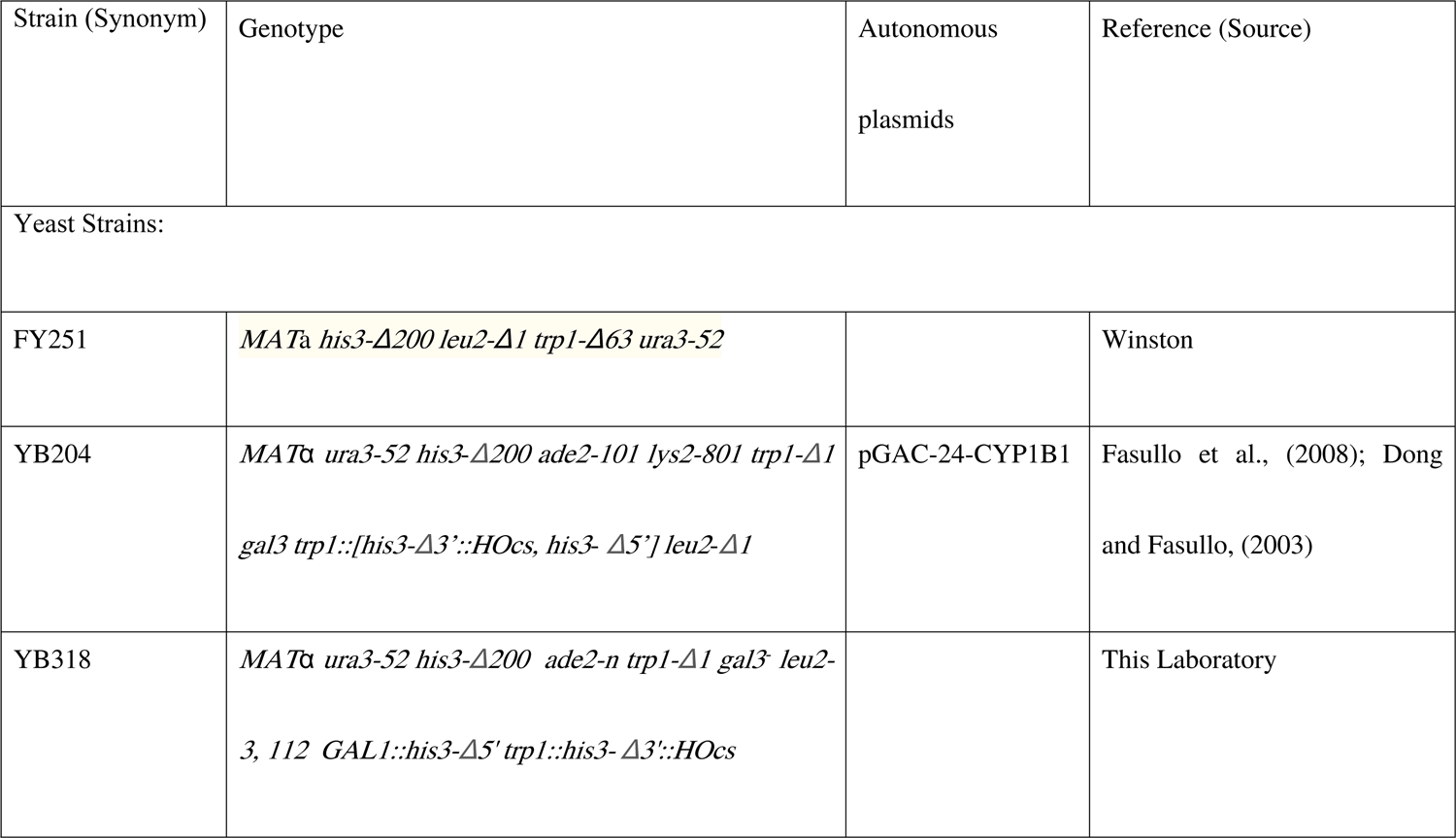
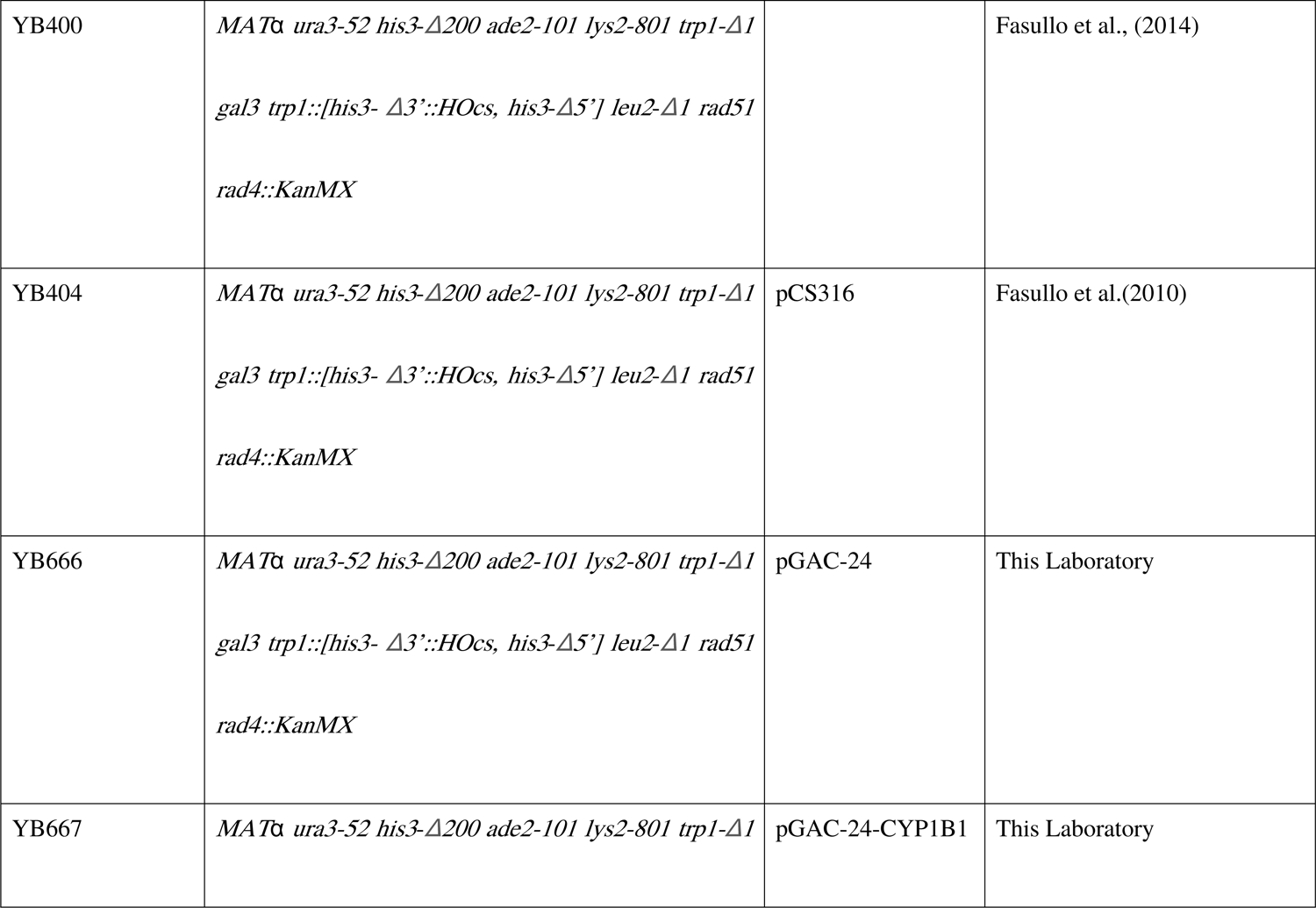
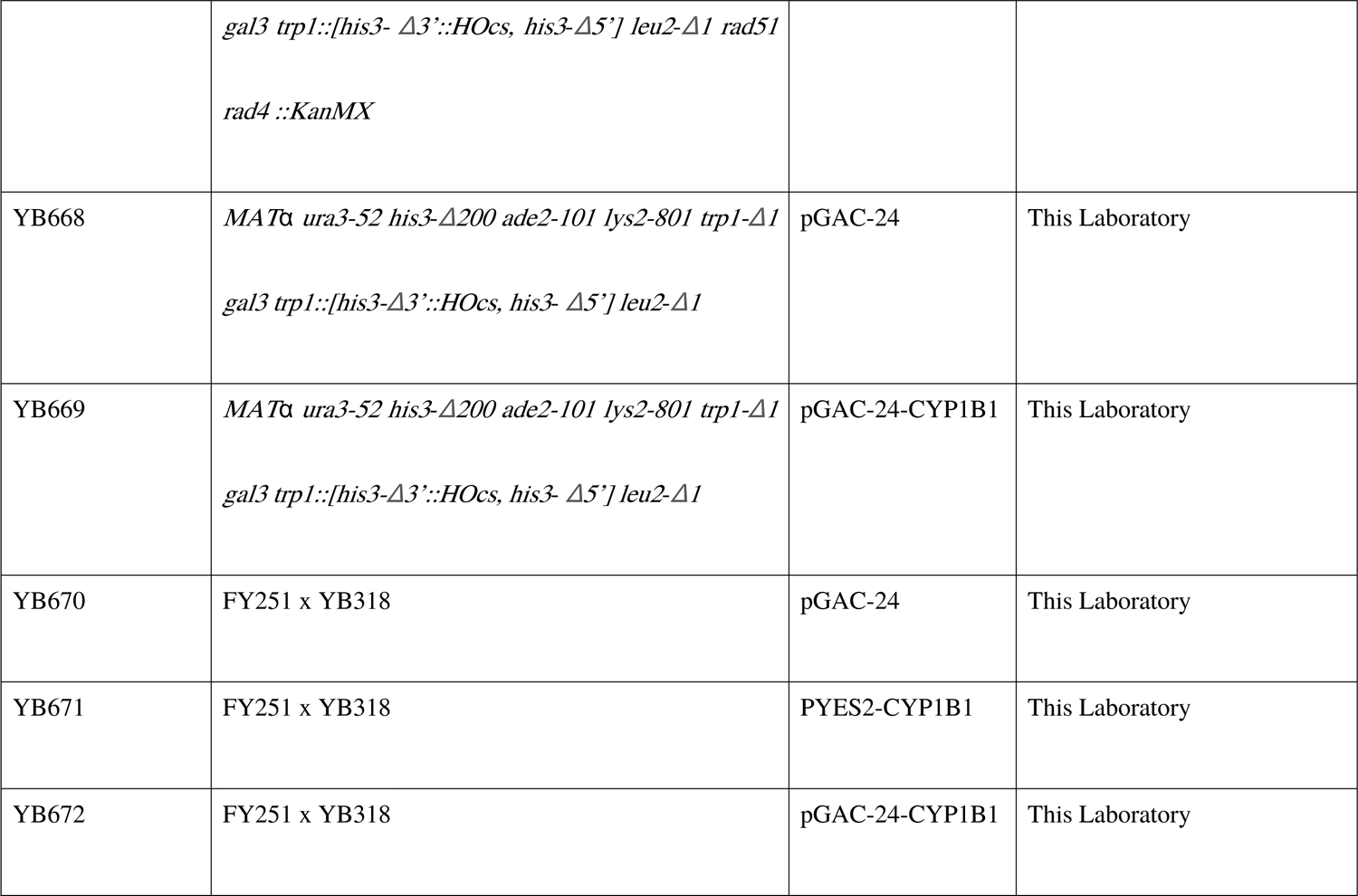
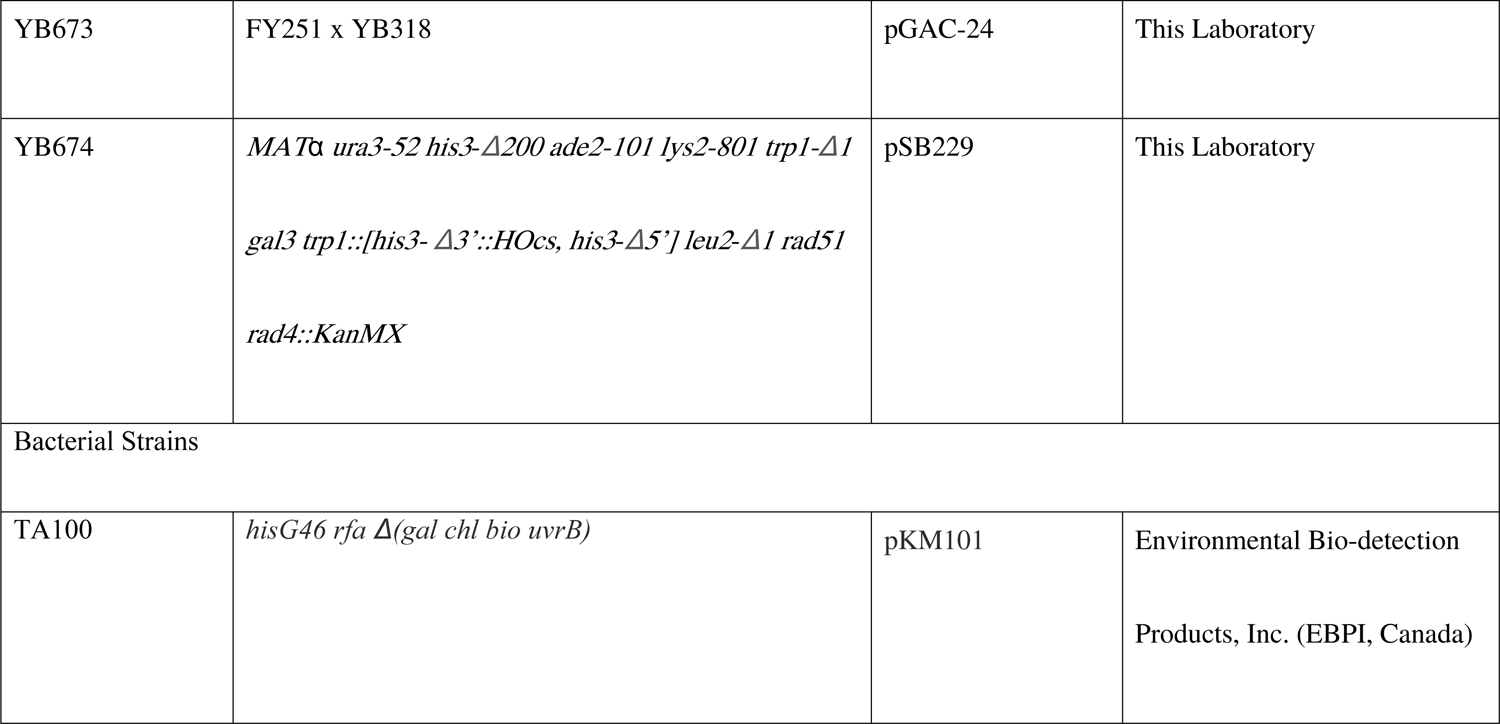
**Yeast and Bacterial Strains**

Monoclonal antibody to CYP1B1 was purchased from Santa Cruz Biochemicals (sc-374228). AFB_1_ was purchased from Sigma Aldrich, BaP-DHD was purchased from Toronto Chemicals, and 2-amino-3-methylimidazo [4,5-f] quinoline (IQ) and 2-amino-3,8-dimethylimidazo[4,5-f] quinoxaline (MeIQx) were purchased from Santa Cruz. The polyaromatic hydrocarbons, AFB_1_ and BaP-DHD, were dissolved in dimethyl sulfoxide (DMSO), while the heterocyclic aromatic amines, IQ and MeIQx, were dissolved in methanol containing 0.1% acetic acid.

### Plasmid Constructions

We originally obtained CYP1B1 from pYES2-CYP1B1, courtesy of T. Sutter (U. Tennessee). In this vector, the CYP1B1 is under the control of the *GAL1* promoter and the *CYC1* terminator. A *Bam*H1-*Bgl*II fragment of CYP1B1 was obtained from pYES2-CYP1B1. pGAC24 contains a polylinker between GPD promoter and PGK terminator (Lesser and Guthrie, 1993). The *Sal*I site was converted to a *Bam*H1 site by commercially available *Bam*H1 linkers. The *Bam*H1-*Bgl*II fragment of CYP1B1 was then subcloned into the new *Bam*H1 site. The presence of the CYP1B1 gene was confirmed by PCR using the forward primer 5’GTCATGAGTGCCGTGTGTTT3’ and reverse primer 5’ATTGGCAAGTTTCCTTGGCT3’, which are located within the coding region of the ORF. The amplification condition was follows: 35 cycles of 30 s at 94°C, 30 s at 52°C, 60 s at 72°C, and a final extension of 10 min at 72°C.The PCR products were then analyzed on a 1% agarose gel.

### Enzyme Assays

CYP1B1 and CYP1A2 enzymatic activity was measured in yeast microsomes by quantifying 7-methoxyresorufin O-demethylase (MROD) (Fasullo et al., 2014), and ethoxyresorufin O-deethylase (EROD) activity (Sengstag et al., 1994). Yeast microsomes were prepared according to Pompon et al. (1996). The enzyme buffer contained 50 mM Tris pH 7.4, 5 μM methoxyresorufin (Sigma) and 500 μM NADPH. Resorufin production was measured in real-time by fluorescence in a Tecan plate reader, calibrated at 535 nm for excitation and 585 nm for absorption, and standardized using serial dilutions of resorufin. The reaction was started by the addition of NADPH and resorufin was measured at one minute intervals during the one hour incubation at 37°C. Rat liver microsomes (S9) were used as a positive control. Enzyme activities were measured in triplicate for at least two independent lysates from each strain and expressed in pmol/min/mg protein.

### Western Blots

Strains were grown to log growth phase (A_600_ = 0.5–1) in SC-LEU at 30°C at 200 rpm. Cells were concentrated and prepared as previously described by Foiani et al. (1994). Samples were preheated at 95°C for 5 minutes with fresh sample buffer (30 mM Tris, 1% SDS, 1% glycerol, 0.002% bromophenol blue, 5% β-mercaptoethanol, pH 6.8). Proteins were then resolved on a 10% SDS-PAGE gel at 100V constant voltage in Tris-Glycine running buffer (25 mM Tris, 192 mM glycine, pH 8.8) for 150 min. Proteins were then transferred to a 0.45 μm PVDF membrane at a constant voltage of 15 V for 35 min in Dunn-Carbonate buffer (10 mM NaHCO_3_, 3 mM Na_2_CO_3_, 20% Methanol, pH 9.9). After transfer, the membrane was blocked in 20 mM Tris 150 mM NaCl Tween 20 0.1% (w/v) (TBST) containing 5% nonfat dry milk for 30 minutes at room temperature at 100 rpm. The membrane was then incubated overnight with primary antibody against CYP1B1 (sc-374228) at 1:1000 in 5% nonfat dry milk in TBST at 4°C with constant shaking. The membrane was then washed three times with TBST (10 minutes each wash) and incubated in secondary antibody (goat anti-mouse IgG-HRP:sc-2005) at 1:10,000 in 5% nonfat dry milk in TBST for 2 hours at room temperature, and washed three times with TBST (10 minutes each wash). Signal was detected via chemiluminescence, using a FluorChem E system. A purified CYP1B1 protein (Creative Biomart) was purchased as a standard in Western blots.

### Growth Curve Assay

In brief, individual saturated cultures were prepared for each yeast strain in selective SC-LEU medium. Cell density was adjusted to ∼0.8 × 10^7^ cells/ml for all cultures. In each microtiter well, 90 μl of YPD medium and 10 μl of cells (8 × 10^4^ cells) were aliquoted in duplicate for blank, control and experimental samples. Cells were exposed to 1 μM, 5 μM, and 10 μM AFB_1_, 1 μM, 5 μM, and 15 μM BaP-DHD, 600 μM and 1 mM IQ, and 750 μM, 1.5 mM, and 3 mM MeIQx.

The microtiter dish was placed in a plate reader that is capable of both agitating and incubating the plate at 30^°^C, as previously described. We measured the A_600_ at 10-minute intervals, for a total period for 24 hours, 145 readings. Data at 1-hour intervals were then plotted. To avoid evaporation during the incubation, the microtiter dishes were sealed with clear optical tape. Area under the curve was determined using freely available software (https://www.padowan.dk/download/).

### Measuring DNA Damage-Associated Recombination and Mutation

To measure carcinogen-associated genotoxic events, log phase yeast cells (A600 = 0.5–1) were concentrated ten-fold in phosphate-buffered saline (PBS) and exposed to indicated doses of AFB_1_ or BaP-DHD, dissolved in DMSO. After exposure, cells were washed twice in PBS, and an undiluted aliquot was inoculated on SC-HIS to select for His^+^ recombinants or SC-ARG + CAN to select for Can^R^. An appropriate dilution was inoculated on YPD to measure viability (Keller-Seitz et al., 2004). Generally, 100–200 colonies are counted to assess viability. Statistical significance was determined by Students t-test (Zar, 1996).

### Electrophoretic Karyotype of His^+^ Recombinants

Intact yeast chromosomes were prepared as previously described (Fasullo and Davis, 1988). Chromosomal DNA was resolved using a CHEF Mapper XA Chiller clamped homogeneous electrical field (CHEF) electrophoresis system (Bio-Rad). The gels were run at a 60° angle for 18 hours with pulse times of 40–75 seconds and then for 8 hours with pulse times of 110–130 seconds. Chromosomal DNA was detected by fluorescence after ethidium bromide staining.

### Inducing CYP1B1 expression

Strains containing CYP1B1 were first inoculated in selective media (SC-URA for pYES2-CYP1B). Cells were then diluted in YP-raffinose. At A_600_ of 0.5-1, galactose or glucose was added to induce or repress the CYP1B1 gene for two hours, and then were exposed to either 50 μM AFB_1_ or 35 μM BaP-DHD in for three hours at 30 °C. After exposure, cells were washed twice in phosphate-buffered saline and plated directly on SC-HIS (glucose) plates to measure recombination and the appropriate dilution was plated on YPD to measure viability.

### Measuring *RNR3* Induction by Rnr3-GFP Fluorescence

*RNR3* induction was measured by green fluorescent protein (GFP) fluorescence. Cells expressing CYP1B1 were inoculated in SC-LEU medium. Log phase cells (A_600_ =0.5–1) were exposed to 100 μM AFB_1_ for 3 h and 24 h, and washed twice in PBS, and then concentrated tenfold. As controls, we measured Rnr3-GFP expression after exposure to 0.05% methyl methanesulfonate (MMS) or in carcinogen-exposed cells not expressing CYP1B1. GFP fluorescence was compared to cells exposed to carcinogen or exposed to the solvent, DMSO, alone. Exposed cells were then counted in an Amnis ImageStream^X^; 1 x 10^4^ cells were analyzed for each sample. Cell shape and GFP fluorescence was measured with both a visible light and a 488 nM laser, using the IDEAS v6.2 software to calibrate cell shape and fluorescence.

### Ames Assay

Cultures of *Salmonella typhimurium* LT2 strains TA100 (*hisG46)*, containing plasmid pKM101, were inoculated in 5 ml of LB-AMP and incubated at 37°C for 16 hours (Mortelmans, 2000). The bacteria were centrifuged and resuspended in 1 mL of LB-AMP (∼2 x 10^9^ cells/mL). The 470 μL master S9-mix, as described by Maron and Ames (1983), was added to 20 μL of rat liver S-9 and 100 μL of the *S. typhimurium* strains; controls substituted sterile H_2_O for rat-liver S9. Tubes were pre-incubated at 37° with defined concentrations of AFB1 or BaP-DHD for 30 minutes. After the pre-incubation, 2 mL of heated top agar (45°C) were added and the contents were poured on minimal glucose plates and incubated at 37°C for 48 hours. His^+^ colonies were then counted.

## RESULTS

### CYP1B1 Expression and Protein Can Be Detected in Budding Yeast

The cDNA of human CYP1B1 had been previously inserted into a yeast expression pYES2, under the control of the inducible GAL promoter (Shimada et al., 1999). To constitutively express CYP1B1, we inserted the cDNA copy of CYP1B1 obtained from pYES2-CYP1B1 in the pGAC-24 (Lesser and Guthrie, 1993) vector under the control of the glyceraldehyde 3-phosphate dehydrogenase (GPD) promoter and the phosphoglycerol kinase (PGK) terminator (Lesser and Guthrie, 1993). To confirm CYP1B1 expression in yeast, we performed 1) Western blots and 2) EROD and MROD activity assays (Table II). Microsomal extracts were obtained from YB204 and YB318 cells expressing constitutive levels of CYP1B1. Both the MROD and EROD activities from yeast microsomal preparations were comparable to previous results from Shimada et al. (1999) and are higher than those obtained using rat S9 microsomes. In addition, microsomal extracts activated more AFB_1_ or BaP-DHD *in vitro*, as measured by His^+^ mutants in the Ames assay, compared to extracts from non-expressing CYP1B1 cells (P = 0.04, Supplemental Table I). TCA extracts were obtained from a yeast strain expressing galactose-inducible CYP1B1, and by Western blot, a protein of approximately ∼58 kDa was identified in cell lysates (supplemental Figure 1). These results indicate that both CYP1B1 enzyme activities and protein can be detected in yeast cells expressing CYP1B1.

**Figure 1.**
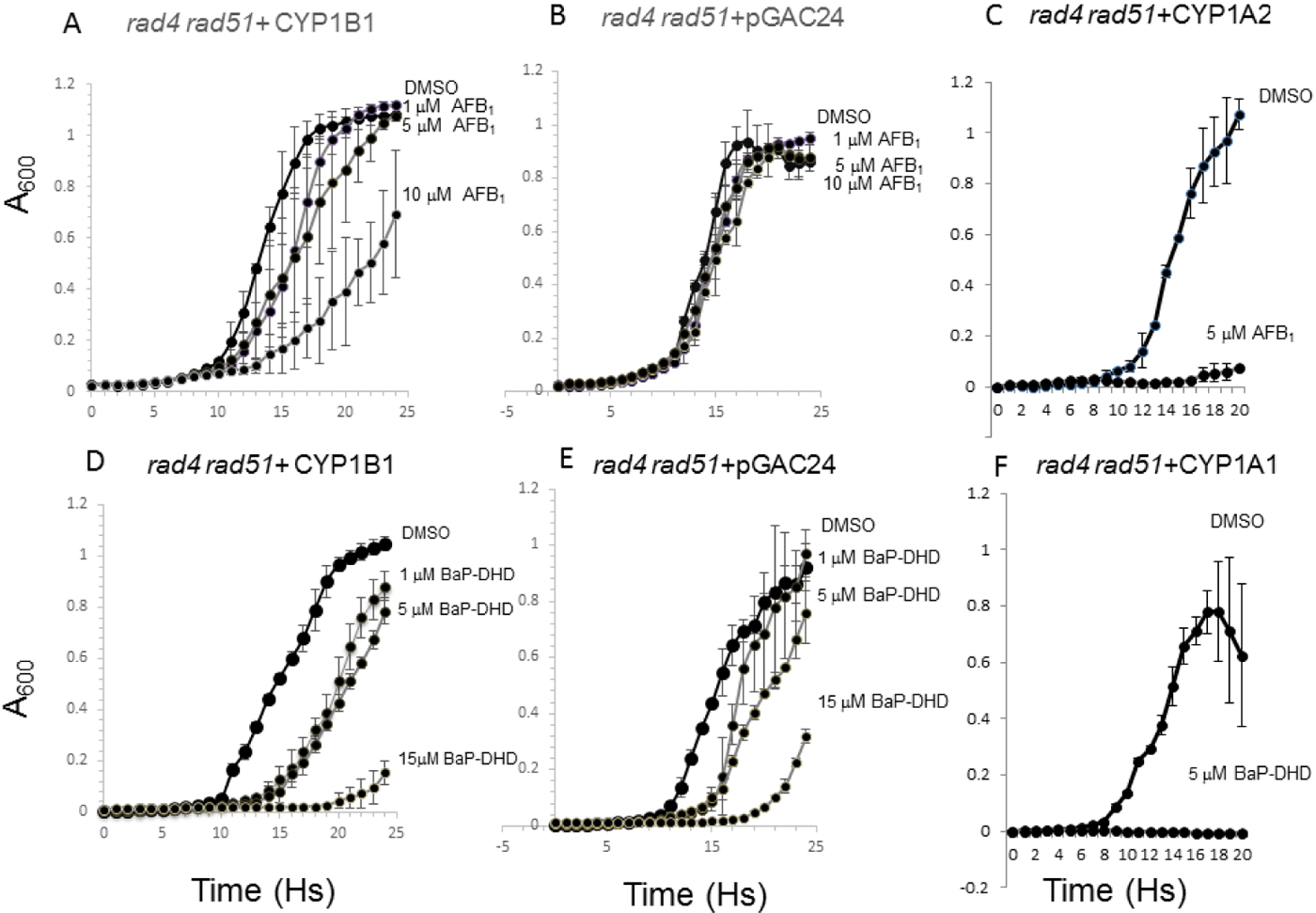
Growth curves of the DNA repair *rad4 rad51* mutant expressing CYP1B1, CYP1A2, CYP1A1 or no CYP (pGAC-24) after exposure to AFB_1_ or BaP-DHD. Approximately 10^5^ cells were inoculated in each well in a 96-well plate platform; yeast growth in each well was monitored in a Tecan Plate reader and over a period of twenty-four hours. Absorbance (A_600_) is plotted against time (h). The first row is growth curves of cells expressing CYP1B1 (A), no CYP (pGAC-24, B), or CYP1A2 (C) after exposure DMSO or indicated doses of AFB_1_. The second row is growth curves of cells expressing CYP1B1 (D), no CYP (pGAC-24, E), or CYP1A1 (F) exposure DMSO or indicated doses of BaP-DHD. Error bars are standard deviations of two biological duplicates. The complete genotype of the *rad4 rad51* mutant is described in Table I.

**TABLE II.**
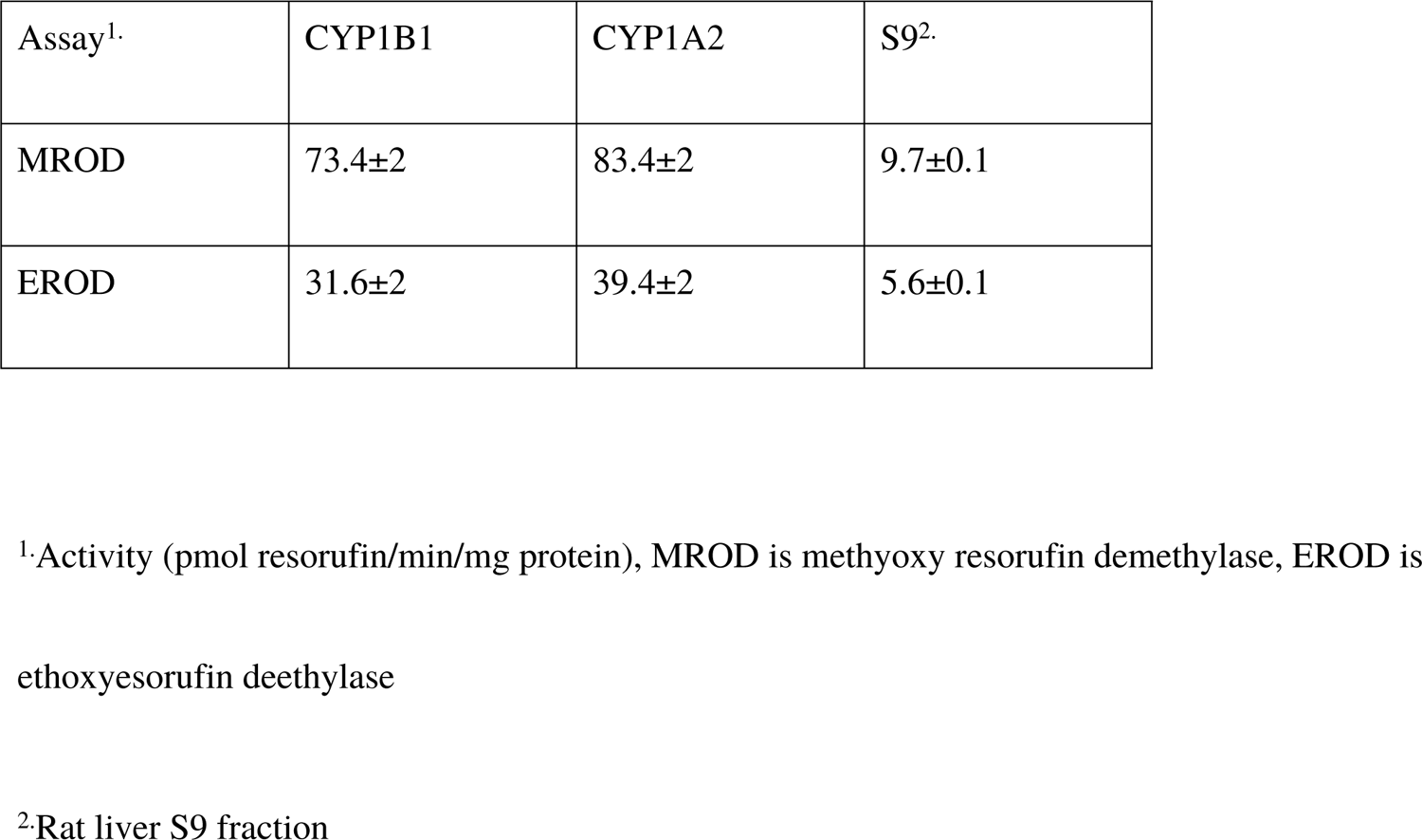
MROD and EROD in yeast (YB204) cells expressing either CYP1B1 or CYP1A2

### Expression of CYP1B1 in Yeast Converts AFB**_1_**, BaP-DHD, IQ, and MeIQx into DNA Damaging Agents

Previously, we showed that expression of human CYP1A1 (Freedland et al., 2017), CYP1A2 (Sengstag et al., 1996; Fasullo et al., 2014), or CYP3A4 (Fasullo et al., 2017) in budding yeast is sufficient to convert polyaromatic hydrocarbons into potent DNA damaging agents. We introduced pGAC-24 (empty vector) and pGAC-24-CYP1B1 in the yeast DNA repair mutant *rad4 rad51*, lacking both nucleotide excision repair and recombinational repair, by selecting for Leu^+^ transformants. By selecting for Ura^+^ transformants, we separately introduced CYP1A1 and CYP1A2, contained on pSB229 and pCS316, respectively, into the *rad4 rad51* strain. Transformants were then exposed to varying concentrations of BaP-DHD and AFB_1_. While expression of CYP1A2 conferred sensitivity to 5 μM AFB_1_, expression of CYP1A1 conferred sensitivity to 5 μM BaP-DHD (Figure 1). Strains expressing CYP1B1 exhibited sensitivity to AFB_1_ in a dose-dependent manner and little sensitivity was observed in the *rad4 rad51* strain containing the empty vector pGAC-24 after exposure to AFB_1_. Likewise, strains containing CYP1B1 exhibited BaP-DHD sensitivity in a dose-dependent manner, although the *rad4 rad51* mutant containing the empty vector did exhibit modest sensitivity to BaP-DHD (Figure 1). Areas under the curve (AUCs) were calculated for growth curves and are shown in supplemental Table II. AUCs calculated for exposed cells containing CYP1B1 are consistently lower than those obtained for exposed cells containing the empty vector. These data show that CYP1B1 expression in yeast is sufficient to confer both AFB_1_ and BaP-DHD sensitivity.

CYP1B1 is expressed in normal colon tissue and in colon tumors (Androutsopoulos et al., 2013) and heterocyclic aromatic amines are colon carcinogens. Expression of CYP1B1 in the *rad4 rad51* DNA repair mutant also conferred modest sensitivity to the heterocyclic aromatic amines IQ and MeIQx (Figure 2). Because both IQ and MeIQx conferred toxicity in strains not expressing CYP1B1, we calculated AUCs and compared them with strains not expressing CYP1B1 (supplemental Table II). Our data indicate that strains that express CY1P1B1 are more sensitive to MeIQx at the highest MeIQx concentration.

**Figure 2.**
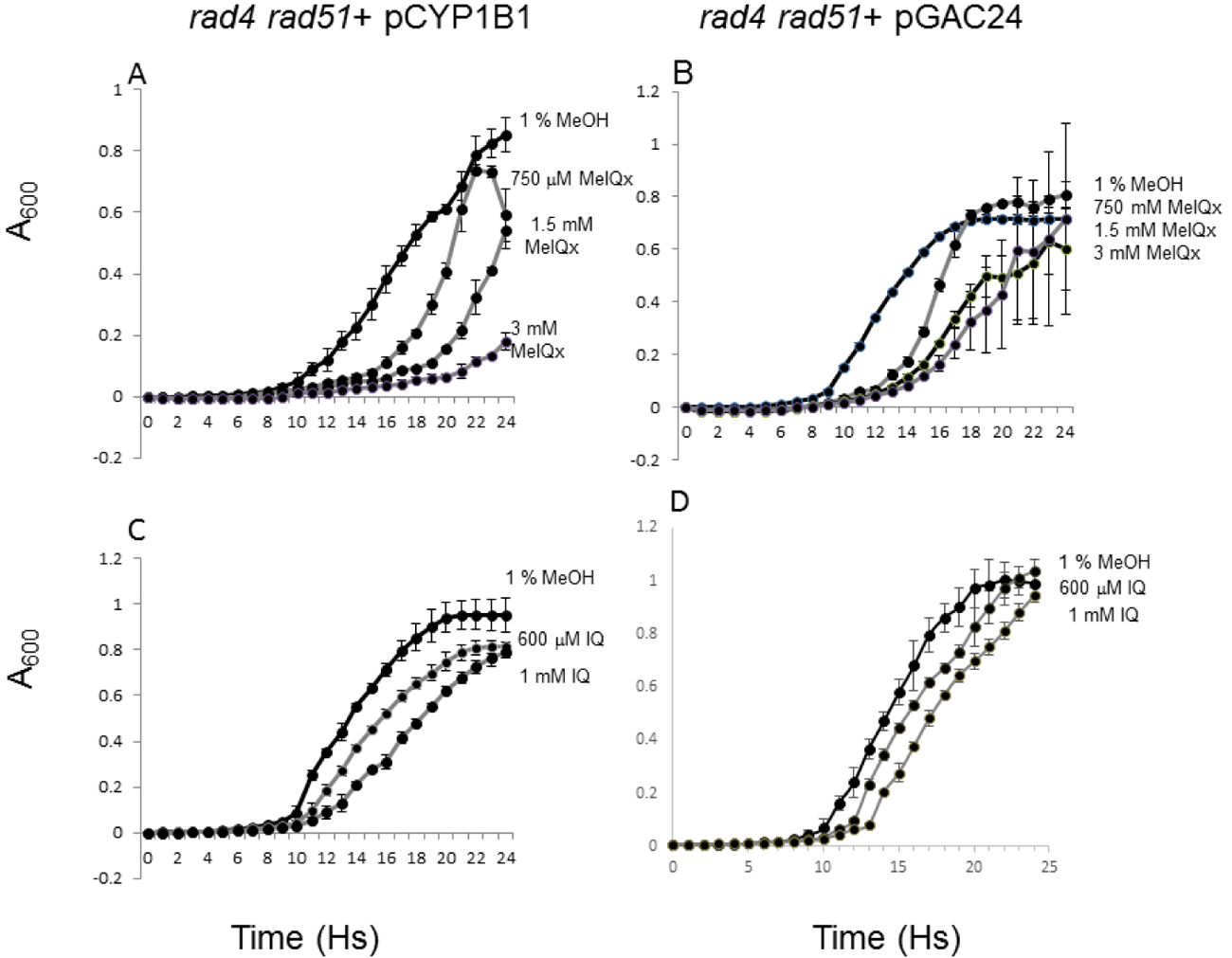
Growth curves of the DNA repair *rad4 rad51* mutant expressing CYP1B1 (YB) or no CYP (pGAC-24) after exposure to MeIQx and IQ. Approximately 10^5^ cells were inoculated in each well in a 96-well plate platform; yeast growth in each well was monitored in a Tecan Plate reader and over a period of twenty-four hours. Absorbance (A_600_) is plotted against time (h). The first row is growth curves of cells expressing CYP1B1 or no CYP (pGAC-24) after exposure MeOH or indicated doses of MeIQx. The second row is growth curves of cells expressing CYP1B1 or no CYP (pGAC-24) after exposure MeOH or indicated doses of MeIQx. Error bars are standard deviations of two biological duplicates. The complete genotype of the *rad4 rad51* mutant is described in Table I.

### CYP1B1 Expression in Yeast Converts AFB**_1_** and BaP-DHD into Recombinogens

We had previously shown that P450-activation of AFB_1_ and BaP-DHD can convert AFB_1_ and BaP-DHD into potent recombinagens and mutagens, respectively. We introduced pYES2-CYP1B1, in which CYP1B1 is inducible by galactose, into a diploid strain that can measure homology-directed ectopic recombination between two *his3* fragments (Figure 3). We exposed CYP1B1-expressing strain AFB_1_ to solvent alone under in galactose medium (YPGal) or glucose medium (YPD). We observed that cells grown in YPGal cells exhibited an approximately eight-fold higher frequency of AFB_1_-associated translocation frequencies compared to cells grown in or that were exposed to solvent alone (Figure 4). Cells grown in glucose, in which CYP1B1 expression is repressed, did not exhibit higher levels of AFB_1_-associated recombination; these data thus revealed that inducible expression was sufficient to stimulate recombination.

**Figure 3:**
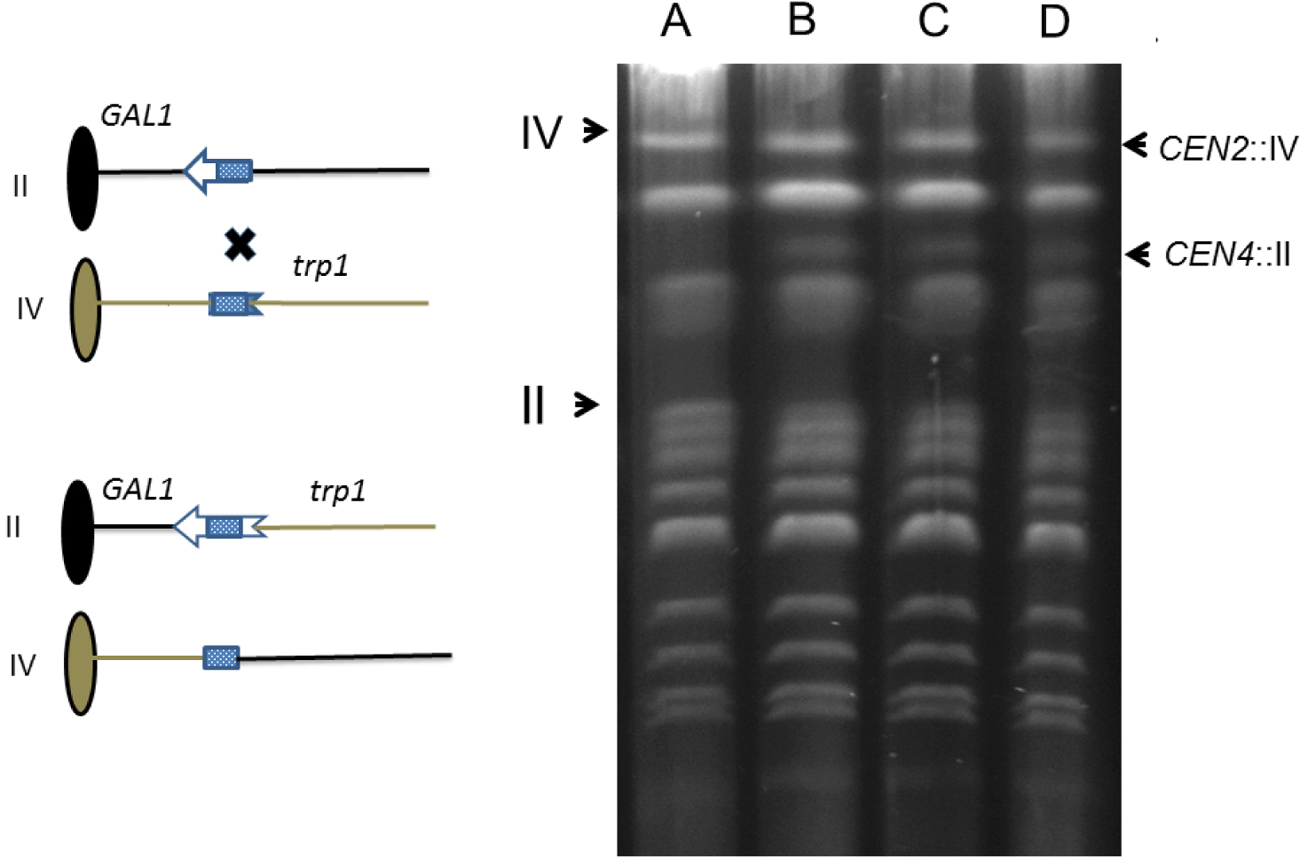
Configuration of *his3* fragments in yeast strains to measure translocations. Figure on the left is a model of the recombination assay and the figure on the right is a pulse field gel of chromosomal DNA obtained from AFB_1_-associated His^+^ recombinants. The oval represents a centromere and the line represents the chromosome; the left arm of the chromosome is not shown for simplicity. The *his3* fragment is shown with arrow and feathers. The similar shaded areas represent shared homology; the dotted pattern represent 446 bp of shared DNA sequence. An “X” denotes where a cross-over event would occur. The product of the recombination event is shown below where *CEN2* is linked to the long arm of chromosome IV (*CEN2*::IV) and *CEN4* is linked to the long arm of chromosome II (*CEN4*::II). The translocated chromosome *CEN4*::II is identified as a novel band on the pulse field gel, while the translocated chromosome *CEN2*::IV, IV, and XII often comigrate. Lane A is the His^-^ parent; lanes B, C, and D are His^+^ recombinants. Chromosomal sizes indicated on the left are based on New England Biolab yeast chromosome pulse field standards.

**Figure 4.**
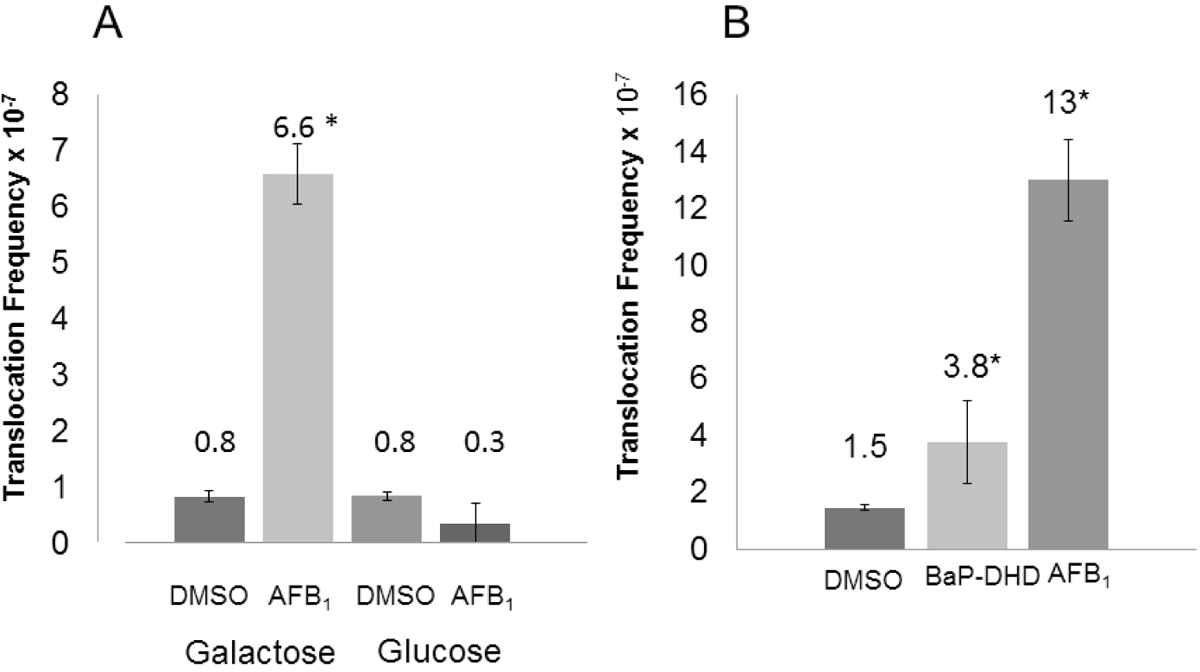
Frequencies of homology-directed translocations in cells expressing either galactose-inducible CYP1B1 (Left) or constitutive CYP1B1 (Right). Approximately 10^8^ cells on SC-HIS plates to select for His^+^ recombinants, and the appropriate dilution was plated on YPD to measure viability (Left). The recombination frequencies are represented in a bar graph for exposure to AFB_1_ (50 μM) or DMSO (1%). CYP1B1 was expressed in cells containing pYES2-CYP1B1 in galactose or glucose medium. The recombination frequencies are listed above the bar graph (Right). Recombination frequencies are represented for cells containing pGAC-24-CYP1B1 after cells were exposed DMSO (1%), AFB_1_ (50 μM), or BaP-DHD (33 μM),. The recombination frequencies are listed above the bar graph. Significant levels of AFB_1_-associated recombination were observed after cells were exposed to 50 μM AFB_1_ as determined by Students t-test, and an asterisk indicates (P < 0.05), compared to the untreated control

After exposure to either BaP-DHD or AFB_1_, cells expressing a constitutive level of CYP1B1 exhibited a threefold or eight-fold higher recombination frequencies, respectively, compared to frequencies obtained after exposure to solvent (DMSO) (P < 0.05). In both of these studies the survival percentage after exposure was ∼50%, compared to solvent alone. The fold stimulation in AFB_1_-associated frequencies in cells expressing inducible CYP1B1 were similar to that observed in cells expressing constitutive CYPIB1. These studies illustrate that CYP1B1 expression from either an inducible or a constitutive promoter is sufficient to activate AFB_1_ to become a potent recombinagen in yeast.

In order to confirm that the carcinogen-associated His^+^ recombinants contain reciprocal translocations, we characterized their electrophoretic karyotype. Translocated chromosomes have novel electrophoretic mobility as shown in Figure 3. Among three AFB_1_-associated recombinants, all three contain reciprocal recombination (Figure 3). In addition, the band representing chromosome II is fainter, supporting the notion that one copy of chromosome II has undergone a rearrangement. These data demonstrate that CYB1B1 can activate AFB_1_ and BaP-DHD into recombinagens.

We observed that expression of CYP1B1 in yeast also converted AFB_1_ into a weak mutagen. We introduced CYP1B1 into a strain YB204 and measured frequencies of canavanine resistant mutants (Can^R^) after exposure to 50 μM of AFB_1_, 33 μM BaP-DHD and 1% DMSO. The average frequency of spontaneous Can^R^ mutants was 1.7 x 10^-6^, consistent with previous results, while the average frequency of AFB_1_-associated Can^R^ mutants increased over two-fold (3.7 x 10^-6^, P< 0.05). Frequencies of BaP-DHD-associated Can^R^ mutants increased less than two-fold compared to frequencies of spontaneous Can^R^ mutants, which was not significantly different (P > 0.05). Cells that contain the empty vector (pGAC-24) did not exhibit any increase in mutation frequency after exposure to AFB_1_. These studies indicate that CYP1B1 expression in yeast can increase carcinogen-associated mutations, depending on the carcinogen.

### Rnr3-GFP Expression Is Induced in AFB**_1_**-exposed Cells Expressing CYP1B1

*RNR3* encodes a large regulatory subunit of ribonucleotide reductase that is DNA damage-inducible but whose basal expression is minimal (Elledge and Davis, 1990). We obtained a BY4741-derived strain containing *RNR3*-GFP. Rnr3-GFP expression can be readily detected within 3 hs after cells are exposed to 0.01% MMS. The plasmid pGAC-24-CYP1B1 was introduced into the BY4741-derived strain by selecting for Leu^+^ transformants. The CYP1B1-expressing cells were then exposed to MMS, AFB_1_, BaP-DHD, or DMSO (Figure 5). The percent of cells expressing GFP was then measured after 3 hs, using the Amnis Image Stream. While GFP-expressing cells were detected in 5% of DMSO-exposed cells, GFP-expressing cells were detected in 33% of MMS-exposed cells and 12% of AFB_1_-exposed cells. After a 20-hour exposure we detected GFP in 57% of the MMS-exposed cells, 39% of the AFB_1_-exposed cells (n =4) and 8% of the DMSO-expressed cells. 10% of the BaP-DHD cells expressed GFP, a value significantly above background (P < 0.05), but less than twice above background. GFP florescence was only detected in 7% of the AFB_1_-exposed cells (n=3) not expressing CYP1B1. These data indicate that CYP1B1-associated AFB_1_ metabolites can trigger a DNA damage response in yeast.

**Figure 5.**
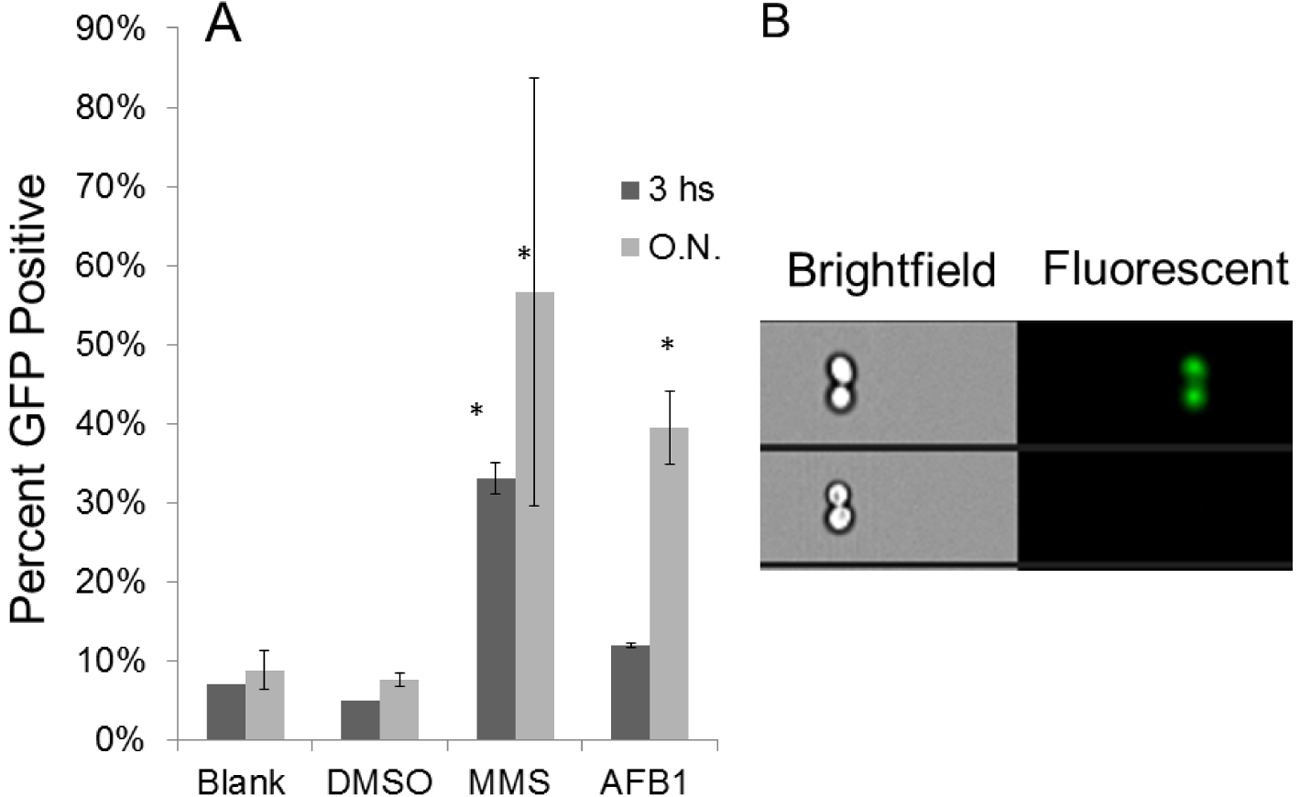
DNA damage-induced expression of *RNR3*-GFP in CYP1B1-expressing yeast cells. A). The percentage of GFP cells is shown on bar graphs after 3 h and 20 h exposure (overnight, ON). The cells were exposed to 0.01% MMS, 100 μM AFB_1_, 1% DMSO, or to no agent. Approximately 10^4^ cells were analyzed for each sample. B). The bright field and fluorescent images of single cells expressing GFP (top) or not expressing GFP (bottom).

## DISCUSSION

CYP1B1 plays a major role in the activation of steroid hormones and xenobiotics. In this manuscript we showed that expression of CYP1B1 in budding yeast was sufficient to activate polyaromatic hydrocarbons and heterocyclic aromatic amines to become DNA damaging agents. In addition, we compared the genotoxicity of BaP-DHD and AFB_1_ in stimulating recombination and mutation. While the activity of CYP1B1 to activate compounds into potent mutagens is well established, our results show that CYP1B1 can activate AFB_1_ and BaP-DHD into recombinagens that can increase frequencies of chromosomal translocations. Furthermore, CYP1B1-activation of AFB_1_ provoked a DNA damage response. Our studies suggest that CYP1B1 expression in higher eukaryotes may be contributing to multiple genetic instability phenotypes that could contribute to carcinogenesis.

We used both mutation and recombination endpoints to measure genotoxicity of CYP1B1-associated metabolites of AFB_1_ and BaP-DHD. Inducible expression of CYP1B1 followed by carcinogen exposure yielded similar AFB_1_-associated recombination frequencies as constitutive expression of CYP1B1 followed by exposure, suggesting that AFB_1_ could be quickly activated in YPGal medium. While frequencies of AFB_1_ and BaP-DHD-associated translocations were significantly above frequencies of spontaneous recombination, higher levels of DNA damage-associated mutations were only significant for cells exposed to AFB_1_. Higher carcinogen-associated frequencies of translocation were easier to detect due to the low frequencies of spontaneous translocation events. Thus, it is possible small increases in BaP-DHD associated mutation frequencies were not detected in this study.

The results are consistent with previous observations that AFB_1_ can be converted into a more potent recombinagen than BaP-DHD (Sengstag et al., 1996, Keller-Seitz et al., 2004). AFB_1_-associated DNA adducts interfere with DNA replication and retard S phase (Fasullo et al. 2010, Lin et al., 2014); whereas BaP-DHD-associated DNA adducts can be effectively bypassed by translesion DNA polymerases (Ikeda et al., 2011). We previously observed that activated BaP-DHD can generate N^2^-Guanine lesions in cells expressing CYP1A1 (unpublished), and CYP1A1-mediated activation of BaP-DHD is strongly mutagenic (Sengstag et al., 1996, Keller-Seitz et al., 2004). Future studies will focus on using NER defective strains to measure CYP1B1 activation of mutagens.

While the *in vitro* EROD and MROD activities of yeast microsomal fractions of CYP1B1 were similar to that of CYP1A2, CYP1B1 expression in yeast cells conferred less carcinogen-associated genotoxicity than CYP1A2 expression. This was true in either strains that expressed or did not express the human oxidoreductase; however, yeast contains an endogenous oxidoreductase (Yabusaki et al, 1988) that may enhance activation of HAAs and polyaromatic compounds (Sengstag et al., 1994). We suggest that observed differences between CYP1B1-mediated and CYP1A2-mediated activation of AFB_1_ are due to subtle differences in protein structure. While the overall volume of the substrate binding cavity is similar, the orientation of the bound substrate differs between CYP1A2 and CYP1B1, likely due to differences in amino acids localized to the edge of the cavity and the protein backbone (Sansen et al., 2007). Two residues in CYP1B1, V395 and A133, are pivotal in the divergent phenotypes. CYP1B1’s V395 is suspected to provide more space for substrate binding than its CYP1A2 residue counterpart, L382 (Wang et al.,2011). Thus, these subtle variations may explain the differences in preferential binding profile observed in these CYPs.

The observation that CYP1B1 can activate HAAs and AFB_1_ in yeast yields insights into immunotoxicity of these compounds (Bellarmi, 201; Gong et al., 2016). While some investigators have suggested the AFB_1_-associated immunotoxicity results from interference with mitochondrial respiration and oxidative stress, CYP1B1 is also induced in human monocytes by AFB_1_ exposure. CYP1B1 expression in the spleen is also suspected to play a role in toxin-associated immunosuppression, and has been observed in mice after exposure to DMBA. Thus, it is possible that AFB_1_ is also a potent genotoxin in monocytes after CYP1B1-mediated activation.

In summary, we have shown that CYP1B1 expression in yeast is sufficient to activate carcinogens into active genotoxins. The yeast expression system will be useful in determining whether cancer-associated CYP1B1 variants confer higher levels of carcinogen genotoxicity.

## AUTHOR CONTRIBUTIONS

Michael Fasullo designed the study. Akaash Kannan performed the Western blots and recombination experiments using pGAC-24-CYP1B1. Nicholas Perpetua performed the growth curves and the recombination experiments using pYES2-CYP1B1 and CYP1A1. Michael Dolan produced and analyzed the Amnis data. All the authors read and approved the study.

## ACKNOWLEDGEMENTS

We thank T. Sutter (U. Tennessee) for pYES2-CYP1B1 and T. Begley for the BY4741 strain containing *RNR3*-GFP. We thank A. Smith and A. Britton for their technical support in the initial stages of this project. This research was supported by grants R21ES015954 and R15ES023685-01 (MF) from the National Institutes of Health.

---

**Supplemental Figure 1:**
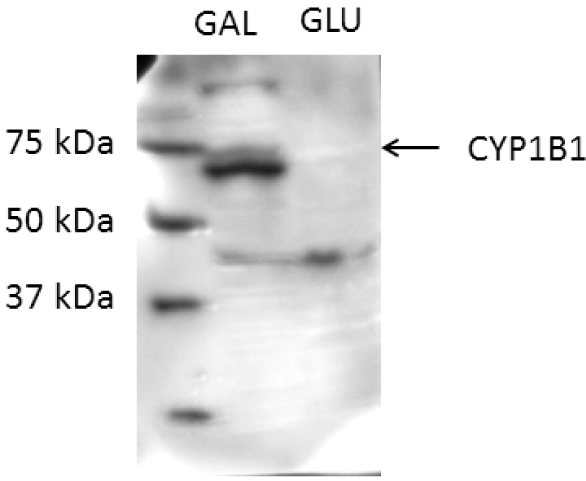
Western blot of cell lysate obtained from cells expressing an inducible CYP1B1. Cells were grown in YP-GAL. The molecular weight markers are shown in the first lane and the protein is shown in the second lane. A band of approximately 58 kDa was detected, as indicated by the arrow.

**SUPPLEMENTAL TABLE I.**
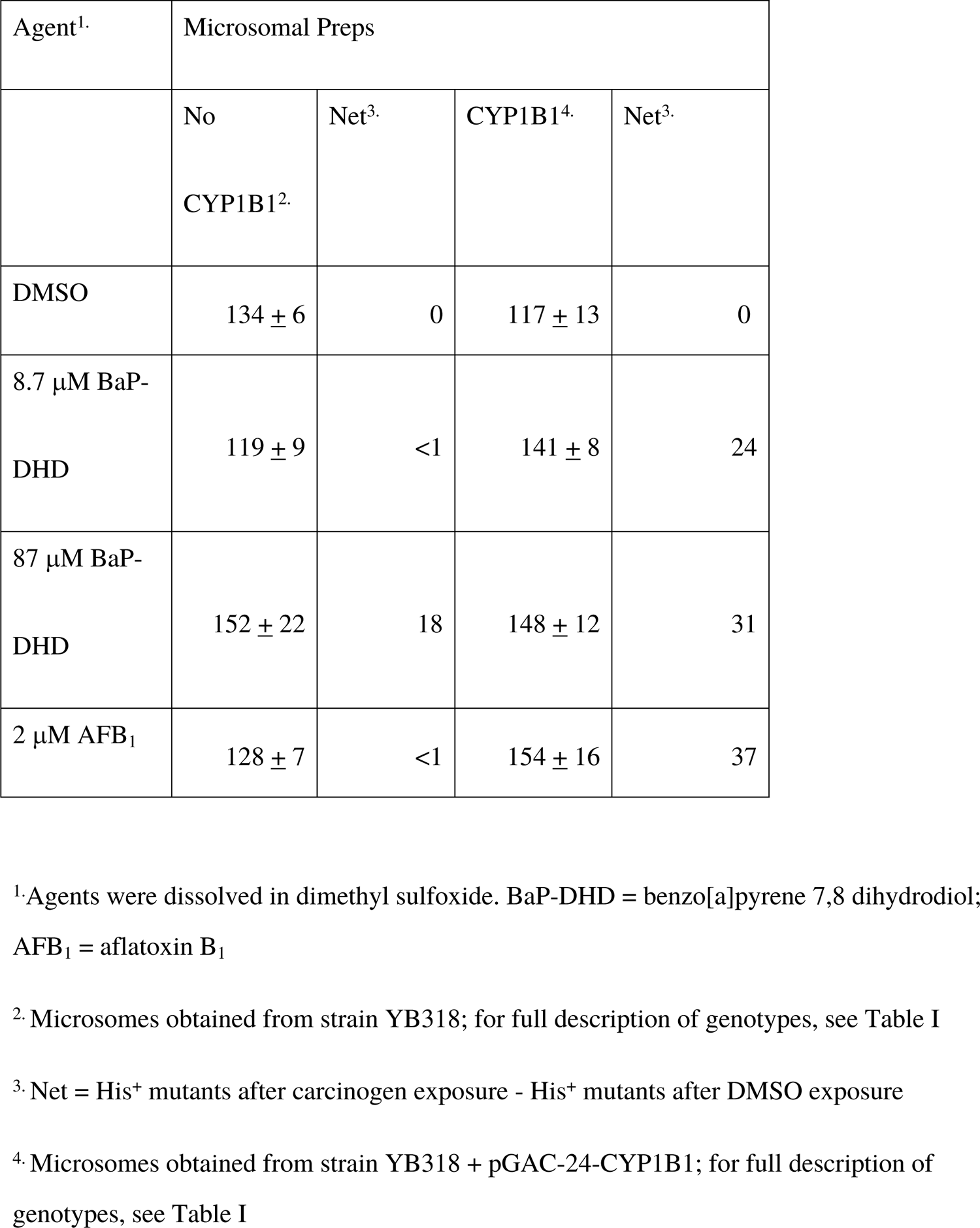
Carcinogen-associated His^+^ Mutants Obtained in the Ames Assay (TA100)

**SUPPLEMENTAL TABLE II.**
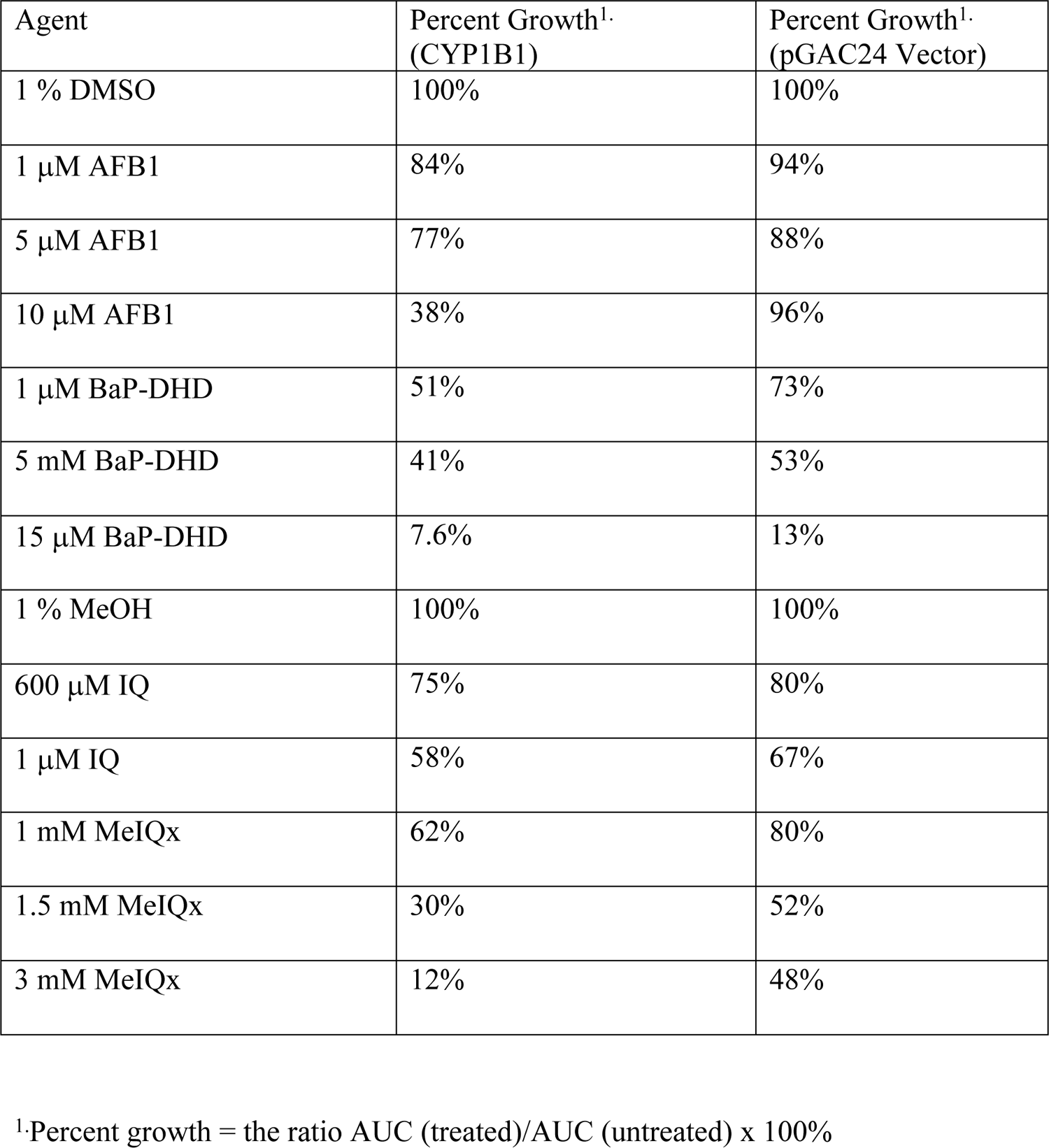
Percent Growth Based on AUCs Obtained from Growth Curves

