## Supplemental Table 1 for "CYP1B1 converts procarcinogens into genotoxins in *Saccharomyces cerevisiae*"

Supplemental Table 1. Carcinogen-associated His^+^ Mutants Obtained in the Ames Assay (TA100)

| Agent^1^ | Microsomal Preps | | | |
| --- | --- | --- | --- | --- |
|  | No CYP1B1^2^ | Net^3^ | CYP1B1^4.^ | Net^3^ |
| DMSO | 134 ± 6 | 0 | 117 ± 13 | 0 |
| 8.7 μM BaP-DHD | 119 ± 9 | <1 | 141 ± 8 | 24 |
| 87 μM BaP-DHD | 152 ± 22 | 18 | 148 ± 12 | 31 |
| 2 μM AFB_1_ | 128 ± 7 | <1 | 154 ± 16 | 37 |

^1^Agents were dissolved in dimethyl sulfoxide. BaP-DHD = benzo[a]pyrene 7,8 dihydrodiol; AFB_1_ = aflatoxin B_1_

^2^ Microsomes obtained from strain YB318; for full description of genotypes, see Table 1

^3^ Net = His^+^ mutants after carcinogen exposure – His^+^ mutants after DMSO exposure

^4^ Microsomes obtained from strain YB318 + pGAC-24-CYP1B1; for full description of genotypes, see Table 1
