## Supplemental Table 2 for "CYP1B1 converts procarcinogens into genotoxins in *Saccharomyces cerevisiae*"

Supplemental Table 2. Percent Growth Based on AUCs Obtained from Growth Curves

| Agent | Percent Growth^1.^ (CYP1B1) | Percent Growth^1.^ (pGAC-24 Vector) |
| --- | --- | --- |
| 1 % DMSO | 100% | 100% |
| 1 μM AFB1 | 84% | 94% |
| 5 μM AFB1 | 77% | 88% |
| 10 μM AFB1 | 38% | 96% |
| 1 μM BaP-DHD | 51% | 73% |
| 5 mM BaP-DHD | 41% | 53% |
| 15 μM BaP-DHD | 7.6% | 13% |
| 1 % MeOH | 100% | 100% |
| 600 μM IQ | 75% | 80% |
| 1 μM IQ | 58% | 67% |
| 1 mM MeIQx | 62% | 80% |
| 1.5 mM MeIQx | 30% | 52% |
| 3 mM MeIQx | 12% | 48% |

^1.^Percent growth = the ratio AUC (treated)/AUC (untreated) x 100%
